# A Shifting Immune Landscape: ILC Redistribution and Neutrophil Polarization in Vascular Cognitive Impairment and Dementia (VCID)

**DOI:** 10.64898/2026.08.18.745638

**Authors:** Lei P Wang, Sahar Emami Naeini, Bidhan Bhandari, Logan Rush, Hannah M Rogers, Hesam Khodadadi, Chandramohan Wakade, Jack C Yu, David C Hess, Évila Lopes Salles, Babak Baban

## Abstract

Vascular cognitive impairment and dementia (VCID) is increasingly recognized as a major contributor to cognitive decline; however, the mechanisms through which vascular dysfunction drives innate immune dysregulation remain poorly understood. In this study, we explore the impact of VCID on the cerebral innate immune landscape, focusing on innate lymphoid cells (ILCs) and neutrophils, two key players in neuroinflammation and brain immune homeostasis.

Using a murine model of VCID induced by bilateral common carotid artery stenosis (BCAS) with modifications in C57BL/6 mice, we investigated innate immune cell distribution, polarization, and functional profiles using flow cytometry and immunofluorescence staining. Our findings reveal a compartment-specific shift in ILC populations, with a reduction of ILC2s in the meninges and concurrent expansion in the choroid plexus, accompanied by altered cytokine production. Furthermore, VCID drove a marked shift in neutrophil polarization toward a pro-inflammatory N1-like phenotype in both the meninges and choroid plexus. Critically, immunofluorescence staining of hippocampal brain sections confirmed that activated N1-like neutrophils, characterized by elevated IL-1β and MPO and reduced IL-10, infiltrate the hippocampal parenchyma in VCID, suggesting a spatially progressive innate immune response spanning from CNS border compartments to brain tissue.

These results identify a novel innate immune signature in VCID, compartment-specific ILC redistribution, pro-inflammatory neutrophil polarization at CNS borders, and parenchymal neutrophil infiltration in the hippocampus, which may collectively amplify neuroinflammation and accelerate cognitive decline, identifying potential therapeutic targets for vascular-related dementia.

**Highlights:**

- VCID reduces meningeal ILC2s while driving their concurrent expansion in the choroid plexus
- N1-like neutrophil polarization is significantly enhanced in VCID meninges and choroid plexus
- Activated N1-like neutrophils infiltrate hippocampal parenchyma in VCID
- N1-like neutrophil activation in VCID is characterized by elevated MPO and IL-1β with concurrent loss of anti-inflammatory IL-10
- A spatially progressive innate immune cascade links CNS borders to brain tissue

**Graphic Abstract:** 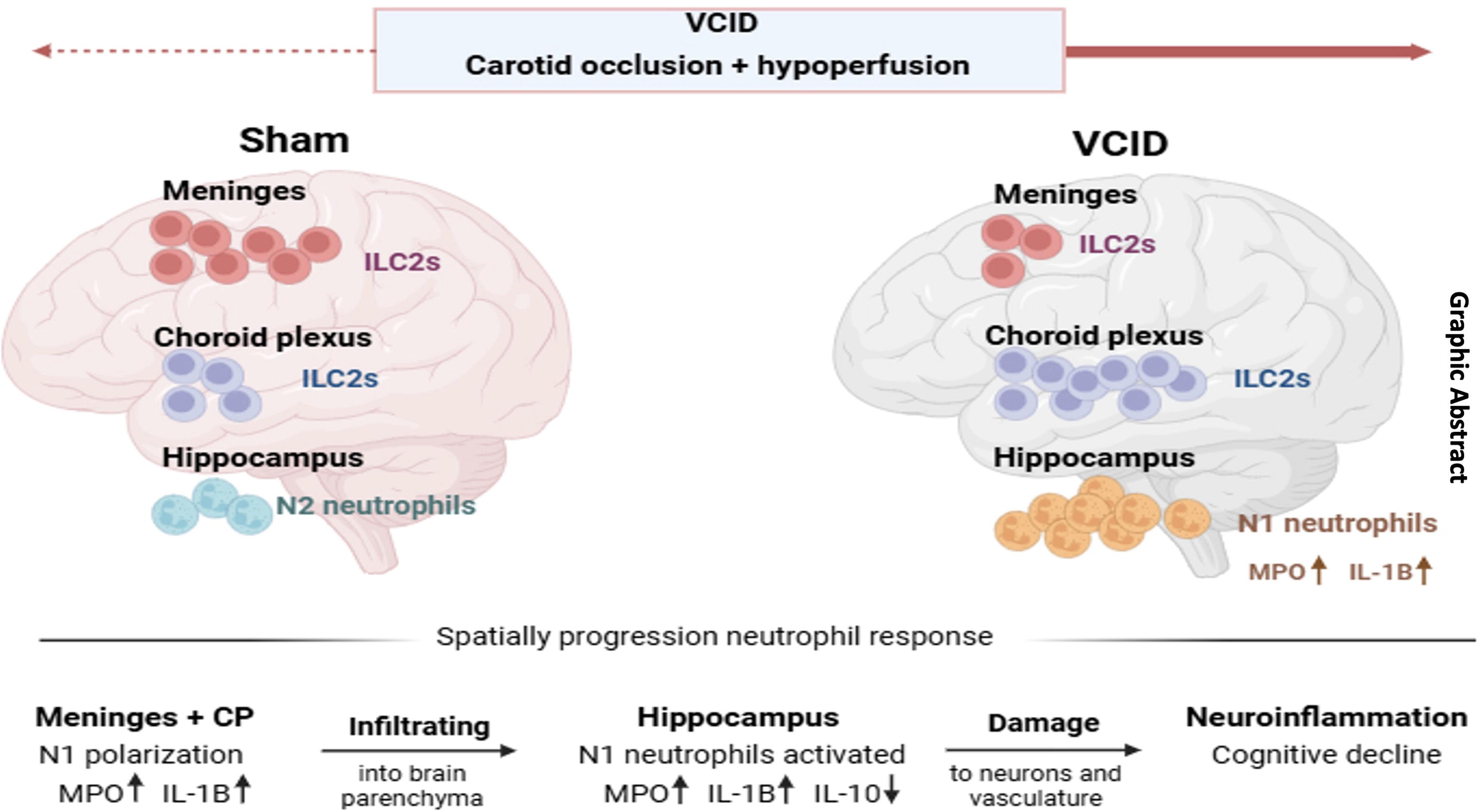

*VCID reshapes the CNS innate immune landscape:* Meningeal ILC2s are significantly reduced while choroid plexus ILC2s expand, reflecting a compartment-specific redistribution. Concurrently, pro-inflammatory N1-like neutrophils, marked by elevated MPO and IL-1β, progressively infiltrate from CNS border compartments into the hippocampal parenchyma, driving neuroinflammation and cognitive decline.

## Introduction

Vascular cognitive impairment and dementia (VCID) includes a spectrum of cognitive disorders from mild deficits to severe dementia (1). Second to Alzheimer’s disease (AD), VCID is potentially responsible for 20% of total dementia cases currently estimated around 36 million, and this number is expected to double by 2030 globally (2). Importantly, several reports indicate the contribution of VCID to the progression of AD by promoting cerebral hypoperfusion, affecting vascular function through endothelial degeneration as well as blood-brain barrier (BBB) dysfunction (3). VCID causes cognitive deficits especially in executive functions and pathology such as white matter disease (WMD), phosphorylated tau (p-tau), amyloid β-protein (Aβ) deposition and neuronal loss. Importantly, despite advances in managing cardiovascular disease and hypertension, there is currently no clear treatment strategy for VCID. The knowledge gap in understanding VCID pathophysiology, particularly mechanisms that connect cerebrovascular defects to neurological and cognitive impairments, remains a critical barrier for developing effective therapy and diagnosis.

The concept of the brain being truly immune privileged has evolved significantly over time (4). While the brain does have unique immunological properties that differ from other organs, it is not truly or completely immune privileged in the absolute sense. Instead, it has a complex and regulated relationship with the immune system that allows for protection of neural tissue while still maintaining necessary immune surveillance and function (4–6).

Under pathologic conditions, vascular and metabolic disarrays may be manifested by cerebral endothelium dysfunction and disintegration of the BBB (7). Such derangement can promote excessive inflammation within the central nervous system (CNS) through local stimulation of inflammatory signals as well as increased infiltration of peripheral immune cells, enhancing further damage and neuroinflammation. Notably, carotid artery stenosis, the primary vascular insult in our VCID model, has been shown to directly damage the blood-CSF barrier and trigger neuroinflammation, further linking vascular dysfunction to CNS immune dysregulation (8).

Aging and cellular senescence can directly contribute to VCID or act as complicating factors in its development (9–10). Inflammaging, the chronic low-grade inflammation associated with aging, disrupts the body’s homeostasis, triggering excessive inflammatory signaling and pro-inflammatory mediator production. This inflammatory environment plays a key role in the onset of VCID and other age-related diseases (11–12). Ultimately, the widespread impact of the CNS and immune system amplifies VCID, transforming it from a localized disturbance to a systemic immunopathological disorder.

The meninges and choroid plexus (CP) are not considered part of the brain tissue itself, but they are integral structures closely associated with the brain that perform essential functions for its protection and support (13–14). Importantly, they serve as critical immunological interfaces between the CNS and the peripheral immune system, regulating immune cell entry, maintaining CNS homeostasis, and orchestrating immune responses to various challenges (15–16). Dysfunction in these structures could potentially contribute to the pathophysiology of VCID through various mechanisms, including altered cerebrospinal fluid (CSF) dynamics, compromised barrier function, and dysregulated immune responses.

Innate Lymphoid Cells (ILCs) represent a relatively recent addition to the immune system, emerging as key architects of immune responses. Like traditional B and T lymphocytes, ILCs originate from lymphoid progenitor cells. However, they differ significantly from conventional lymphocytes due to the absence of recombined antigen receptors and specific lineage markers. These distinctions, along with their ability to respond rapidly to environmental cues, classify ILCs as components of the innate immune system (17–19). ILCs are highly adaptable to the microenvironment of their resident tissues, engaging in dynamic interactions with stromal cells, resident cells, and T cells to maintain homeostasis and protect the host (20–21). In the brain, ILCs are primarily localized in the meninges and CP (22). Recent studies have shown that ILC2s, a key regulatory subset of ILCs, can alleviate symptoms of age-related cognitive decline (23–24), underscoring their potential therapeutic relevance in neurological health. Furthermore, recent evidence from our group demonstrated that adoptively transferred ILC2s can actively traffic to and localize within CNS compartments, including in the context of glioblastoma, further underscoring their unique migratory capacity and therapeutic potential within the brain (25). Given the critical position of the meninges and CP as immunological gateways between the CNS and the periphery, the unique profile and compartmental distribution of ILCs in these regions may represent valuable targets for immunotherapy in VCID. Notably, the impact of VCID on the activation, polarization, and compartmental distribution of ILCs within these CNS interfaces remains largely unexplored.

While ILCs serve as the master regulators and architects of innate immune responses, orchestrating the inflammatory tone and directing the behavior of downstream effector cells, neutrophils represent the frontline foot soldiers of innate immunity. As the most abundant circulating innate immune cells, neutrophils are among the first responders to vascular injury and inflammation (26). Under homeostatic conditions, neutrophil infiltration into the brain is tightly regulated. However, in the context of cerebrovascular damage, BBB disruption, and sustained neuroinflammation, hallmarks of VCID, neutrophils can breach this barrier and accumulate within CNS compartments, where they exert powerful pro- or anti-inflammatory effects depending on their polarization state (27).

Neutrophils are now recognized to adopt distinct functional phenotypes: the pro-inflammatory N1 phenotype, characterized by high expression of myeloperoxidase (MPO), interleukin-1β (IL-1β), and reactive oxygen species (ROS); and the anti-inflammatory N2 phenotype, marked by elevated arginase and IL-10 expression (28). The balance between N1 and N2 neutrophil polarization has profound implications for the resolution or amplification of neuroinflammation. Beyond polarization, activated N1 neutrophils can undergo NETosis, the release of neutrophil extracellular traps (NETs), web-like structures composed of DNA, histones, and granular proteins including MPO, which has been shown to significantly exacerbate neurological damage and worsen outcomes in CNS injury models (29). Whether VCID-associated neutrophil activation similarly drives NETosis and contributes to disease progression represents an important and compelling question for future investigation.

Critically, the crosstalk between ILCs and neutrophils is emerging as a key regulatory axis in innate immunity, ILCs, through their cytokine secretion, can directly influence neutrophil recruitment, activation, and polarization, while neutrophils in turn shape the local inflammatory microenvironment that ILCs inhabit (23). Understanding this ILC-neutrophil axis in the context of VCID, where both vascular damage and sustained neuroinflammation converge, represents a critical and underexplored frontier.

In this study, we sought to advance our understanding of the innate immune landscape in VCID by investigating the compartment-specific distribution and polarization of both ILCs and neutrophils in the meninges and choroid plexus. By elucidating how these two key innate immune populations shift and interact in the context of VCID, we aim to identify novel immune signatures and potential therapeutic targets that could help slow or halt the progression of vascular-related cognitive decline.

## Materials and Methods

### Animals

Wild-type male C57BL/6 mice (n=10 per group, total n=20), 20–24 weeks of age, were obtained from Jackson Laboratories (Bar Harbor, ME) and used to generate the VCID model. Animals were housed in the laboratory animal facilities of Augusta University with free access to food and water. All experiments were performed in accordance with National Institutes of Health (NIH) guidelines and regulations. All experimental procedures were approved by the Institutional Animal Care and Use Committee (IACUC) of Augusta University (protocol No. 2011–0062, renewed February 2026).

### Induction of experimental VCID

Experimental VCID was induced using the bilateral common carotid artery stenosis (BCAS) model as previously described (30). Anesthesia was induced by exposing mice to a gaseous mixture of 30% oxygen, 70% N₂O, and 2.5% isoflurane via vaporizer, and maintained at 1.5% isoflurane throughout the procedure. Mice were kept on spontaneous respiration via breathing mask, positioned in the supine position, and secured to a heating pad with adhesive tape. Body temperature was monitored and maintained at 37 ± 0.5°C throughout the surgical procedure.

A sagittal ventral midline neck incision (∼1 cm) was performed, and both salivary glands were carefully separated and mobilized to expose the underlying common carotid arteries (CCAs). Each CCA was gently isolated from the accompanying vagal nerve and adjacent veins without injuring these structures. Customized steel microcoils (inner diameter 0.18 mm) were applied to both the right and left CCAs at 30-minute intervals to induce bilateral stenosis and establish chronic cerebral hypoperfusion. Normal saline was applied throughout to maintain wound moisture.

Following the procedure, wounds were closed, and lidocaine ointment (Xylocaine 2%; AstraZeneca GmbH, Wedel, Germany) was applied as needed for postoperative analgesia. Mice were transferred to a pre-heated (30°C) recovery box until full consciousness was regained. Sham-operated mice underwent identical surgical procedures with the exception of micro coil placement. Immune profiling was initiated four weeks post-surgery, consistent with the chronic hypoperfusion phase previously characterized using this model (30).

### Immunofluorescence staining

Brain tissues were collected and processed for immunofluorescence imaging as described previously (31). Briefly, coronal brain sections were prepared and incubated with fluorescent-conjugated antibodies targeting neutrophil phenotypic and functional markers, including LY6G, MPO, IL-1β, and IL-10 (all from BioLegend, San Diego, CA, USA), to assess neutrophil infiltration and polarization status within the hippocampal parenchyma of sham and VCID mice. Two separate panels were employed: DAPI/LY6G/IL-1β/MPO to evaluate pro-inflammatory N1-like neutrophil activation, and DAPI/LY6G/IL-1β/IL-10 to assess anti-inflammatory N2-like markers. Hippocampal regions were identified using standard anatomical landmarks and consistent coronal coordinates across all animals. Slides were counterstained with DAPI (4′,6-diamidino-2-phenylindole) to identify viable cell nuclei prior to examination and microscopic imaging. Fluorescence pixel intensity for each marker was quantified using Leica MetaMorph software, and mean pixel intensity values were compared between sham and VCID groups.

### Preparative and analytical flow cytometry

Single cell suspensions from meninges and choroid plexus (CP) were analyzed using a NovoCyte Quanteon flow cytometer (Agilent Technologies, Santa Clara, CA, USA) as described previously (19).

#### ILC Analysis

Cells were gated as Lin⁻CD45⁺ lymphocytes using a lineage depletion cocktail of FITC-conjugated antibodies (all from BioLegend unless otherwise noted) targeting CD3 (clone 17A2), CD4 (clone GK1.5), CD8 (clone 53-5.8), CD14 (clone Sa14-2), CD15 (clone MC-480), CD16 (clone 93), CD19, and CD20 (clone SA271G2). ILC subsets were identified within the Lin⁻CD127⁺ gate as follows: ILC1 (IL-12Rβ2⁺; R&D Systems), ILC2 (GATA3⁺), and ILC3 (RORγt⁺; Thermo Fisher Scientific). Cytokine production was assessed for ILC1s (IFN-γ/TNF-α), ILC2s (IL-5/IL-13), and ILC3s (IL-17/IL-22) as previously described (20). The FlowJo UMAP plugin was used to visualize the distribution and clustering patterns of ILC2s in the meninges and CP before and after VCID (32).

#### Neutrophil Analysis

Total neutrophils were identified as CD45⁺CD11b⁺LY6G⁺ cells. Polarization phenotypes were assessed within the LY6G⁺ gate: N1-like pro-inflammatory neutrophils were defined as LY6G⁺IL-1β⁺MPO⁺, while N2-like anti-inflammatory neutrophils were defined as LY6G⁺IL-10⁺Arginase⁺. The N1/N2 ratio was calculated as an index of neutrophil polarization status. Additionally, the FlowJo UMAP plugin was applied to neutrophil populations from meninges and CP to visualize the distribution and clustering patterns of N1-like and N2-like neutrophils before and after VCID, enabling unbiased assessment of polarization shifts across compartments.

For both ILC and neutrophil panels, isotype-matched controls were used to set appropriate gates, with positivity defined as exceeding a 2% isotypic control threshold. All data were analyzed using FlowJo software (version 10.9; FlowJo LLC, Ashland, OR, USA).

### Statistics

Data analysis was performed using two-way analysis of variance (ANOVA) to assess the effects of treatment (sham vs. VCID) and brain compartment (meninges vs. choroid plexus) on ILC and neutrophil frequencies, polarization states, and cytokine expression levels. Post-hoc pairwise comparisons were conducted using Tukey’s Honest Significant Difference (HSD) test to determine specific group differences following a significant ANOVA result. For immunofluorescence data, pixel intensity values for LY6G, IL-1β, MPO, and IL-10 were compared between sham and VCID groups using unpaired Student’s t-test. The N1/N2 neutrophil ratio was compared between groups and compartments using two-way ANOVA followed by Tukey’s post-hoc test. The assumption of normality was evaluated using the Shapiro-Wilk test, and where violated, non-parametric alternatives including the Mann-Whitney U test or Kruskal-Wallis test were employed. A significance level of p < 0.05 was considered statistically significant. All statistical analyses were performed using SPSS (IBM SPSS Statistics 30).

## Results

### VCID altered the frequencies and polarization of ILCs in the meninges

Flow cytometry analysis revealed a significant reduction in the frequencies of all meningeal ILC subtypes in VCID mice compared to sham controls (Fig. 1). Additionally, VCID induced a notable shift in ILC polarization within the meninges. While ILC3 was the predominant subtype in sham controls, ILC1 emerged as the relatively predominant meningeal ILC subtype following VCID, suggesting a shift toward a pro-inflammatory innate immune tone at this CNS interface.

**Figure 1.**
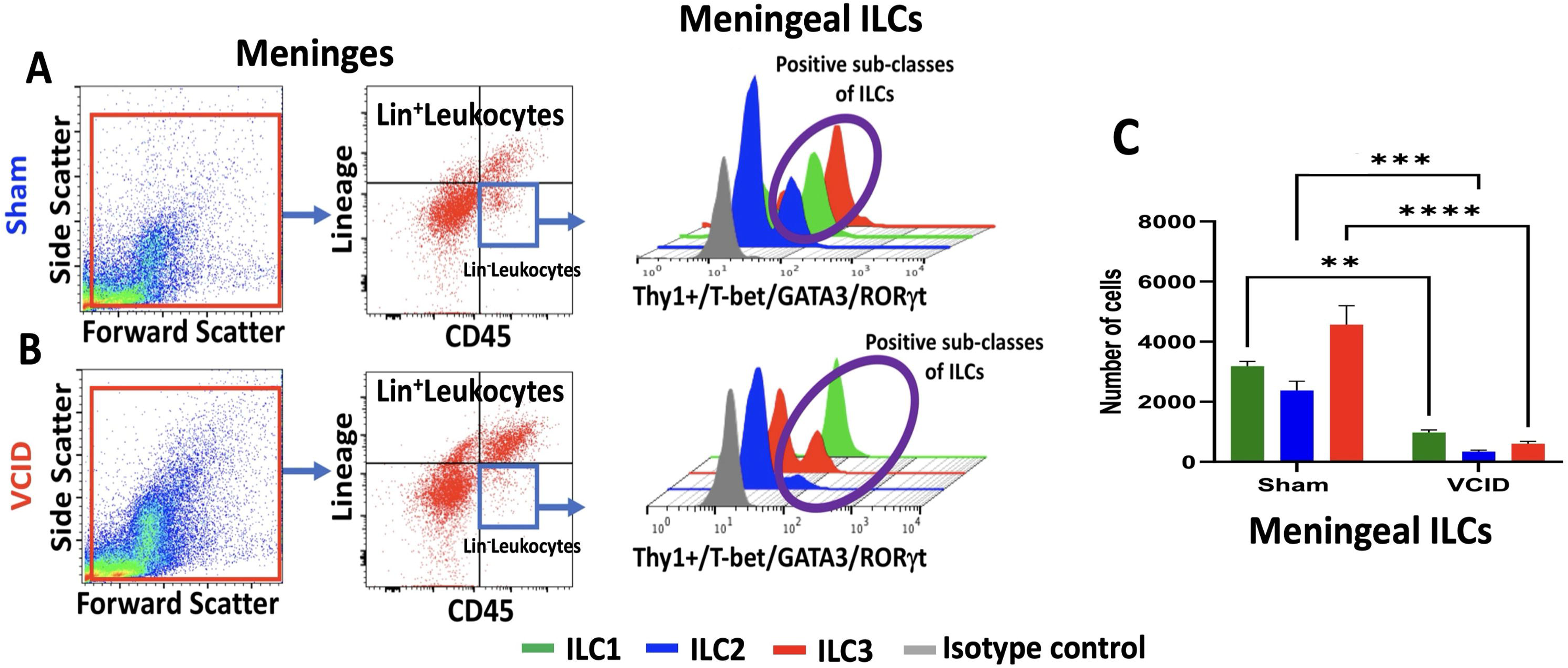
VCID alters the frequencies and subtype distribution of ILCs in the meninges. Flow cytometry analysis of meningeal single-cell suspensions from sham and VCID mice. Panels A and B display the sequential gating strategy, from FSC/SSC scatter to Lin⁻CD45⁺ leukocytes, and the corresponding ILC subtype histograms (ILC1: green; ILC2: blue; ILC3: red; isotype control: gray) for sham and VCID groups, respectively. Panel C presents quantification of meningeal ILC frequencies, demonstrating a significant reduction in all ILC subtypes following VCID compared to sham controls (** p ≤ 0.01, *** p ≤ 0.001, **** p ≤ 0.0001).

### VCID altered the frequencies and polarization of ILCs in the choroid plexus

Flow cytometry analysis revealed a significant increase in total ILC frequency in the choroid plexus (CP) of VCID mice compared to sham controls (Fig. 2A-B). However, this overall increase was not uniformly distributed across all ILC subtypes. As shown in Fig. 3C, ILC2s exhibited the most substantial increase following VCID (*p* < 0.001), followed by ILC1s (*p* < 0.01), while ILC3s showed a significant reduction (*p* < 0.01) compared to sham controls, mirroring the pro-inflammatory shift observed in the meninges, but with a distinct compartment-specific pattern.

**Figure 2.**
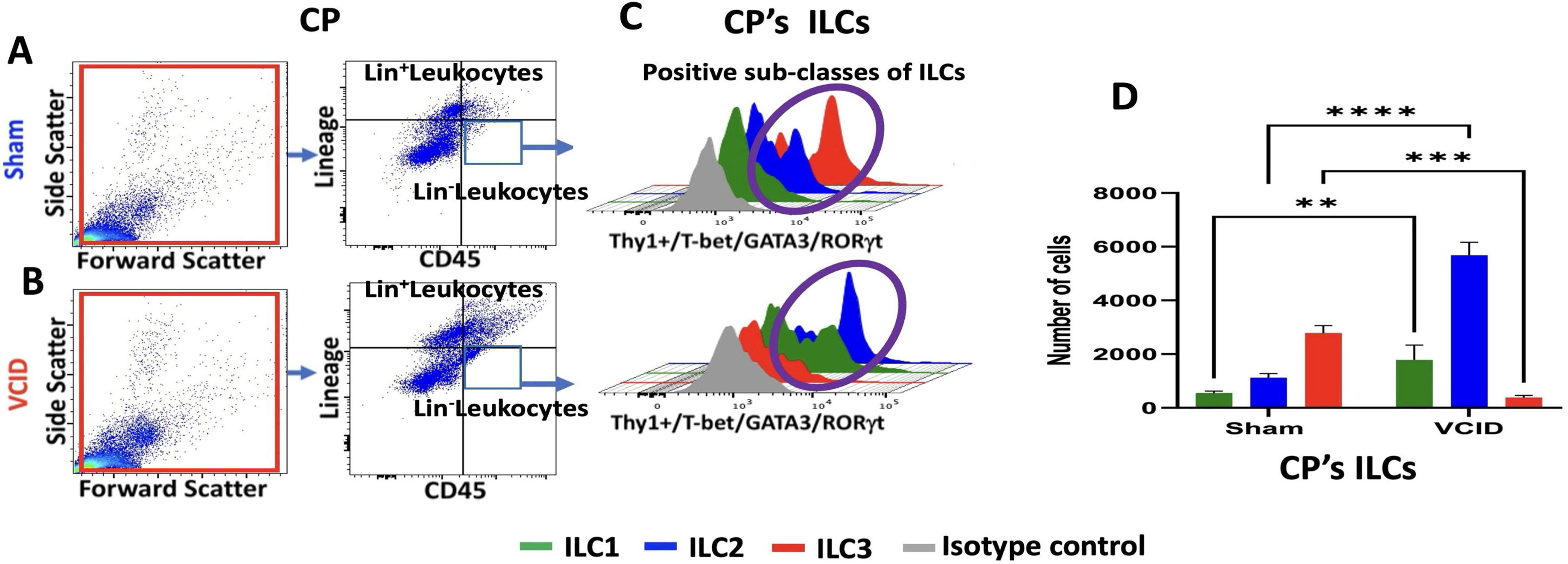
VCID alters the frequencies and subtype distribution of ILCs in the choroid plexus (CP). Panels A and B display the sequential gating strategy, from FSC/SSC scatter to Lin⁻CD45⁺ leukocytes, for sham and VCID groups respectively. Panel C shows the overlay histogram profiles (ILC1: green; ILC2: blue; ILC3: red; isotype control: gray) confirming that VCID leads to increases in ILC1 and ILC2 subpopulations while ILC3 is significantly reduced within the CP. Panel D presents quantification of CP ILC frequencies, demonstrating significant differences between VCID and sham groups (* p ≤ 0.01, *** p ≤ 0.001, **** p ≤ 0.0001).

**Figure 3.**
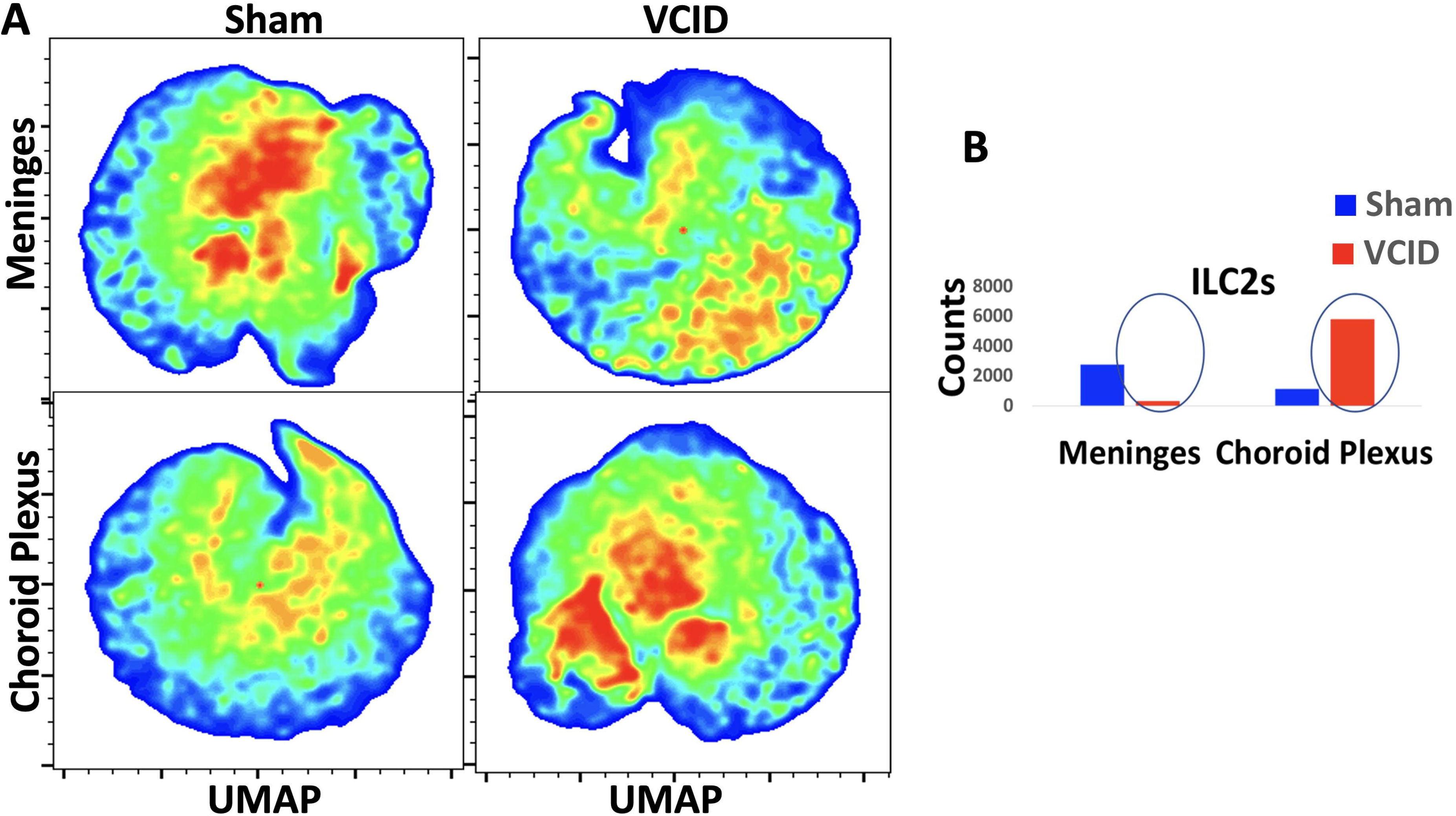
VCID induces compartment-specific redistribution of ILC2s in the meninges and choroid plexus. Panel A displays UMAP density plots generated using the FlowJo UMAP plugin, illustrating changes in the frequency and distribution of ILC2s in the meninges (top row) and choroid plexus (bottom row) of sham and VCID mice. Warm colors (red/orange) indicate high-density ILC2 clusters, while cool colors (blue) represent low-density regions. VCID is associated with a marked reduction in ILC2 density in the meninges and a concurrent expansion of ILC2 clusters in the choroid plexus, consistent with a compartment-specific redistribution. Panel B presents quantification of ILC2 counts in the meninges and choroid plexus, confirming a significant reduction in meningeal ILC2s and a reciprocal increase in choroid plexus ILC2s following VCID (blue: sham; red: VCID).

### VCID shifted the distribution pattern and Functional Features of ILC2s in the Meninges and Choroid Plexus

UMAP analysis revealed significant changes in both the frequency and distribution of ILC2s in the meninges and CP of VCID mice compared to sham controls (Fig. 3). VCID was associated with a reduction in ILC2s in the meninges and a concurrent expansion of the ILC2 population in the CP, a compartment-specific redistribution with important functional implications given the established role of ILC2s in cognitive function (22–23). Flow cytometry analysis further revealed that VCID resulted in a decrease in IL-10 production by meningeal ILC2s, coinciding with their reduced frequency (Fig. 4). In contrast, key ILC2-associated cytokines in the CP, including IL-5, IL-9, and IL-13, were significantly elevated, reflecting the expansion of the ILC2 population in this compartment. Together, these findings suggest that VCID induces a heightened and compartment-specific inflammatory response, with the choroid plexus emerging as a critical site of innate immune dysregulation.

**Figure 4.**
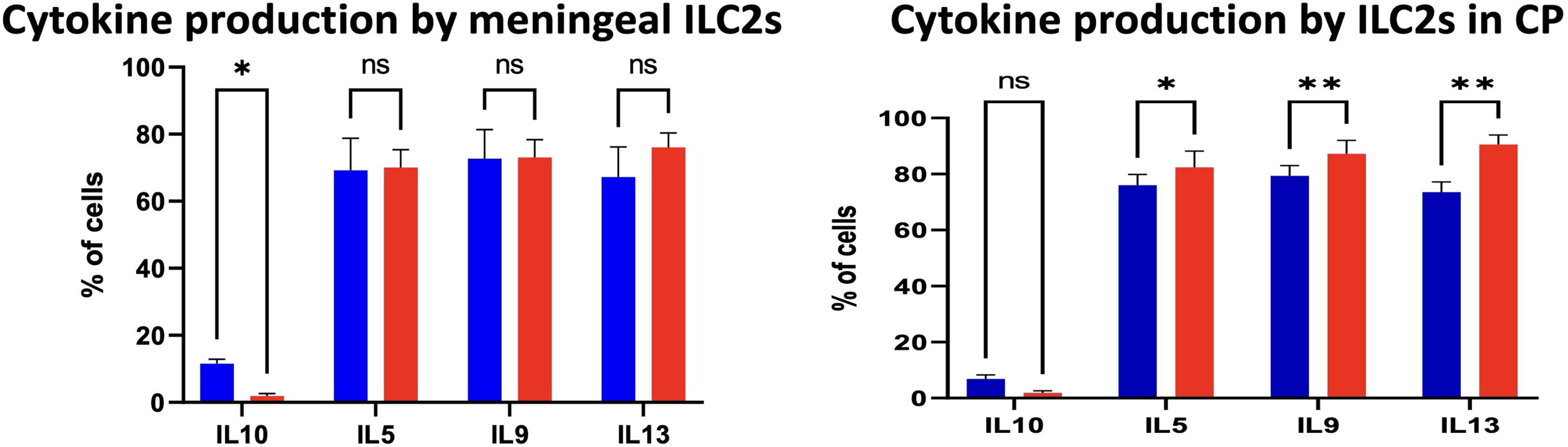
VCID alters cytokine production by ILC2s in the meninges and choroid plexus. Bar graphs depict the percentage of cytokine-producing ILC2s (blue: sham; red: VCID) in the meninges (left panel) and choroid plexus (right panel). In the meninges, VCID significantly reduced IL-10 production by ILC2s (* p ≤ 0.05), while IL-5, IL-9, and IL-13 levels remained unchanged (ns). In the choroid plexus, VCID significantly elevated IL-5 (* p ≤ 0.05), IL-9 (** p ≤ 0.01), and IL-13 (** p ≤ 0.01) production, reflecting the functional expansion of ILC2s in this compartment, while IL-10 showed no significant change (ns). Together, these findings indicate that VCID drives a compartment-specific shift in ILC2 cytokine profiles, with loss of anti-inflammatory IL-10 in the meninges and enhanced type 2 cytokine production in the choroid plexus (ns: non-significant, * p ≤ 0.05, ** p ≤ 0.01).

### VCID Drives Pro-Inflammatory Neutrophil Polarization in the Meninges, Choroid Plexus and Hippocampal Parenchyma

To further characterize the innate immune landscape in VCID, we investigated neutrophil infiltration, activation, and polarization using complementary flow cytometric and immunofluorescence approaches. Flow cytometry analysis revealed a significant increase in the frequency of N1-like pro-inflammatory neutrophils (CD45⁺CD11b⁺LY6G⁺IL-1β⁺MPO⁺) in both the meninges and choroid plexus of VCID mice compared to sham controls, while N2-like anti-inflammatory neutrophils (LY6G⁺IL-10⁺Arginase⁺) were significantly reduced (Fig. 5A). The N1/N2 ratio was markedly elevated in VCID mice in both compartments, indicating a pronounced shift toward a pro-inflammatory neutrophil phenotype (Fig. 5B).

**Figure 5.**
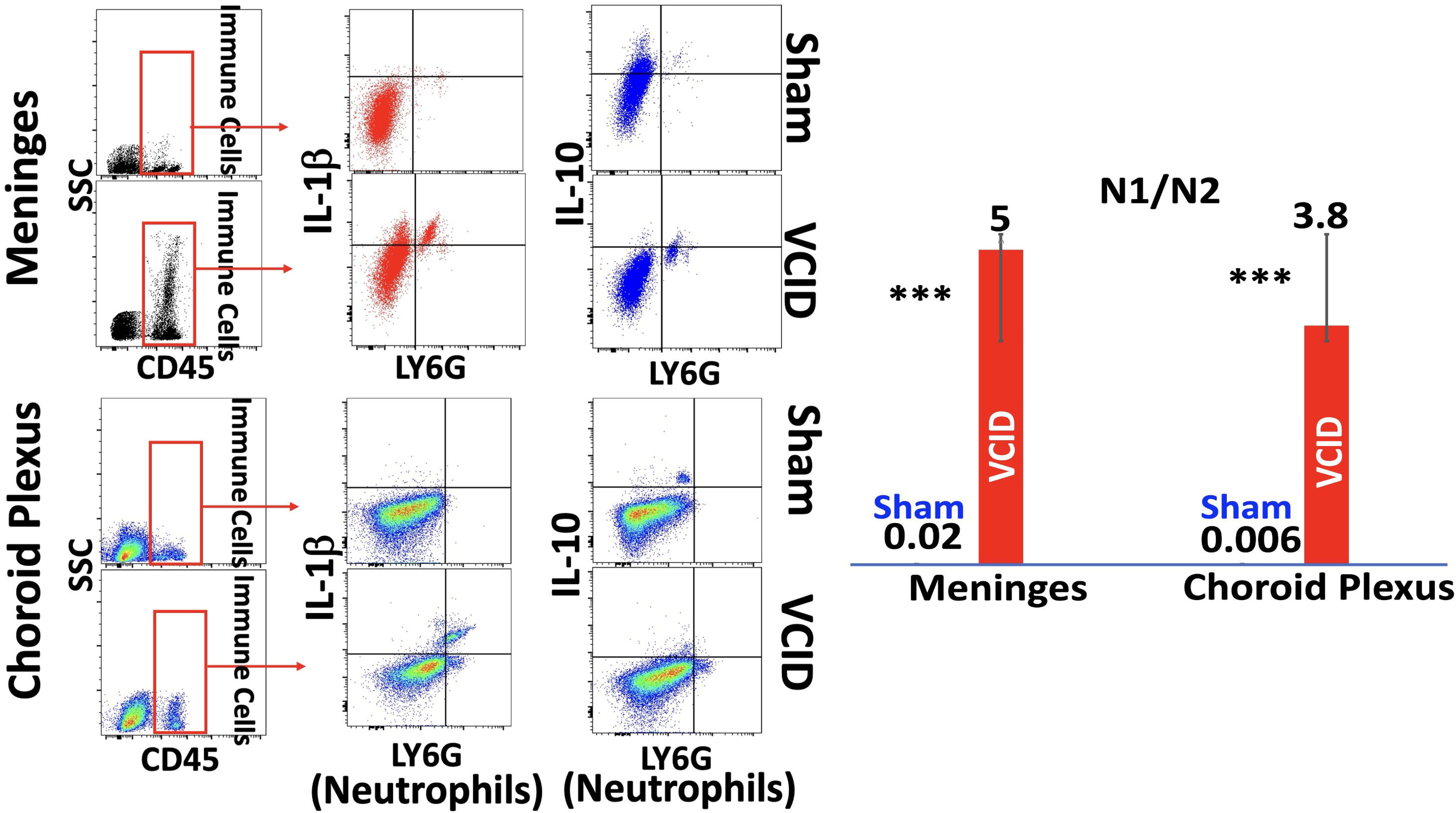
VCID drives a pro-inflammatory shift in neutrophil polarization across CNS border compartments. Flow cytometry analysis of neutrophil polarization was performed in meningeal and choroid plexus single-cell suspensions from sham and VCID mice. Panel A displays the sequential gating strategy — from SSC/CD45⁺ immune cells to LY6G⁺ neutrophils — followed by dot plot analysis of N1-like (LY6G⁺IL-1β⁺) and N2-like (LY6G⁺IL-10⁺) neutrophil populations in the meninges (top row) and choroid plexus (bottom row) for sham and VCID groups. Panel B presents the quantified N1/N2 neutrophil ratio in both compartments, demonstrating that VCID induced a striking pro-inflammatory shift compared to sham controls, with N1/N2 ratios rising from 0.02 to 5.0 in the meninges and from 0.006 to 3.8 in the choroid plexus (*** *p* ≤ 0.001). These findings indicate that under homeostatic conditions neutrophils at CNS borders maintain a predominantly anti-inflammatory N2-like state, which is profoundly disrupted by VCID-induced cerebral hypoperfusion.

To determine whether neutrophil activation at these CNS border compartments is accompanied by parenchymal infiltration and tissue-level inflammatory activity, we performed immunofluorescence staining on hippocampal brain sections. Staining with DAPI/LY6G/IL-1β/MPO (Fig. 6A, top row) demonstrated markedly elevated co-expression of IL-1β and MPO in LY6G⁺ neutrophils within the hippocampus of VCID mice compared to sham controls, confirmed by significantly increased mean pixel intensity for both LY6G and IL-1β (*p* < 0.001; Fig. 6B). This finding confirms that neutrophils not only accumulate and polarize at the meningeal and choroid plexus borders but also infiltrate the hippocampal parenchyma, where their pro-inflammatory N1-like activity is sustained. In contrast, staining with DAPI/LY6G/IL-1β/IL-10 (Fig. 6A, bottom row) revealed a substantial reduction in IL-10 expression in hippocampal neutrophils from VCID mice, consistent with the loss of anti-inflammatory N2-like activity observed in the flow cytometry data (Fig. 6B).

**Figure 6.**
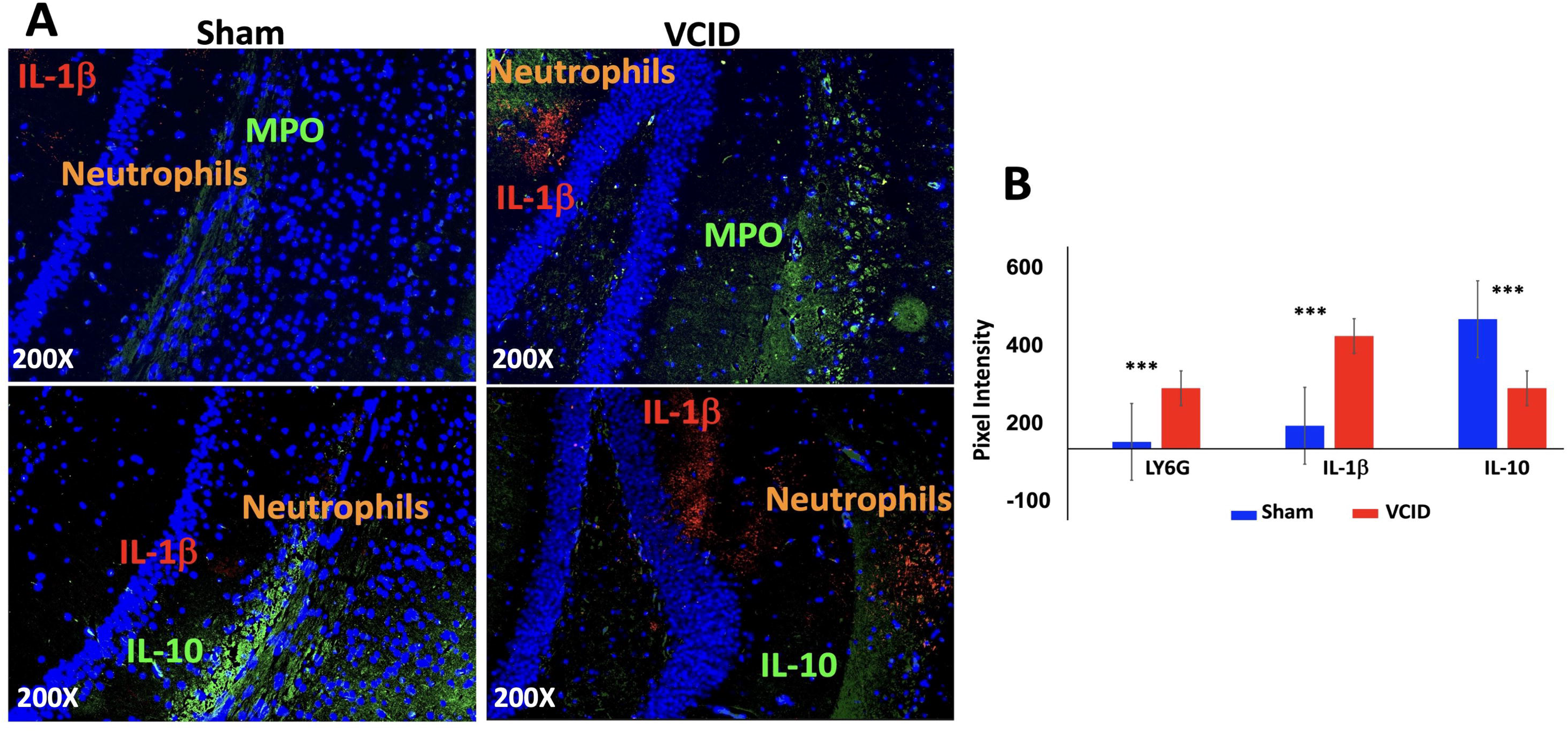
VCID promotes N1-like neutrophil infiltration and pro-inflammatory activation in the hippocampus. Immunofluorescence staining of coronal hippocampal sections from sham and VCID mice. Panel A displays representative multiplex fluorescence images showing neutrophil infiltration and polarization markers across two staining panels: the N1-like panel (top row) co-staining for neutrophils/LY6G (orange), IL-1β (red), and MPO (green); and the N2-like panel (bottom row) co-staining for neutrophils/LY6G (orange), IL-1β (red), and IL-10 (green); nuclei counterstained with DAPI (blue). VCID mice demonstrate markedly increased hippocampal neutrophil infiltration with robust co-expression of IL-1β and MPO, indicative of N1-like pro-inflammatory activation, alongside a concurrent reduction in IL-10 signal, reflecting loss of N2-like anti-inflammatory activity. Panel B presents mean pixel intensity quantification confirming that VCID significantly increased LY6G and IL-1β expression while significantly reducing IL-10 compared to sham controls (*** *p* ≤ 0.001; blue: sham, red: VCID). Together with Figure 6, these findings establish that VCID drives hippocampal neutrophil infiltration with a polarized N1-like phenotype, providing a tissue-level correlate of the CNS border immune dysregulation observed by flow cytometry.

Together, these findings demonstrate a coherent and spatially progressive innate immune response in VCID — neutrophils polarize toward a pro-inflammatory N1-like phenotype at the CNS border compartments of the meninges and choroid plexus, and subsequently infiltrate the hippocampal parenchyma where their sustained activation, characterized by elevated MPO and IL-1β and reduced IL-10, likely contributes directly to neuronal damage and cognitive decline.

## Discussion

Our knowledge about the brain’s immune status has dramatically changed over the past two decades. We now recognize that the brain has a unique and complex relationship with the immune system, involving active regulation and compartmentalized access rather than complete isolation (4). This evolving understanding has important implications for our approach to neurological diseases and potential therapies. Importantly, recognizing that the brain is not an immune-privileged organ significantly enhances our understanding of VCID, highlighting the critical role of neuroinflammation, the importance of BBB function, the impact of systemic inflammation, and opening new possibilities for diagnosis and treatment (7).

ILCs are important and increasingly recognized members of the innate immune system, including within the CNS. Their discovery and characterization have significantly expanded our understanding of innate immunity and its interactions with the brain (17–18). This growing body of knowledge carries significant implications for how we approach neurological diseases and the development of potential therapies.

The novel findings in this study regarding ILCs are significant for two key reasons. First, we confirmed a notable difference in the ILC profile between the meninges and the CP under homeostatic conditions. Second, and more importantly, our data reveal for the first time that VCID alters the profile and distribution pattern of ILCs differentially across these two compartments. Specifically, VCID shifts the balance of ILCs, increasing the proportion of ILC2s in the CP relative to the meninges. Several factors may explain this compartment-specific redistribution. VCID is characterized by significant vascular changes, including hypoxia, chronic ischemia, and endothelial dysfunction, that disrupt normal immune surveillance and compromise BBB integrity (8). These alterations may facilitate the infiltration of inflammatory cells and molecules into brain tissue and influence the polarization and activation of ILCs, driving a shift toward ILC2s in the CP. The altered distribution and activation of ILC2s could contribute to an imbalance in pro- and anti-inflammatory mediators, potentially exacerbating the chronic, low-grade inflammation characteristic of VCID. Although these mechanisms align with our current understanding of ILC2 function in the brain, the precise link between ILC2 redistribution and VCID pathophysiology warrants further investigation. The intricate interactions between immune cells, vascular health, and cognitive outcomes in VCID remain an active and important area of research.

Complementing the ILC findings, our study provides the first evidence of a spatially progressive neutrophil inflammatory response in VCID, spanning from CNS border compartments to brain parenchyma. Flow cytometry analysis demonstrated a significant shift toward pro-inflammatory N1-like neutrophil polarization in both the meninges and choroid plexus of VCID mice, characterized by elevated IL-1β and MPO expression and reduced IL-10 and Arginase levels (27–28). Critically, immunofluorescence staining of hippocampal brain sections revealed that this neutrophil activation is not confined to the CNS borders, LY6G⁺ neutrophils expressing high levels of MPO and IL-1β were detected within the hippocampal parenchyma of VCID mice, with significantly elevated pixel intensity compared to sham controls. This finding suggests that neutrophils activated at the meningeal and choroid plexus interfaces subsequently infiltrate deeper brain structures, bringing their pro-inflammatory activity directly into hippocampal tissue, a region critically vulnerable to vascular injury and neuroinflammation in VCID (33–34).

The elevated MPO expression observed in hippocampal neutrophils is particularly noteworthy. MPO is a key mediator of oxidative stress and tissue damage, and its upregulation in N1-like neutrophils reflects an activated, potentially destructive phenotype (27–28). Furthermore, MPO is a critical component of neutrophil extracellular traps (NETs), web-like structures of DNA, histones, and granular proteins released during NETosis (26). Previous work from our group demonstrated that NETs significantly exacerbate neurological deficits following traumatic brain injury (29), raising the compelling possibility that NETosis may similarly contribute to hippocampal damage and vascular injury in VCID. Whether VCID-associated N1 neutrophil activation progresses to NETosis in the hippocampus, and whether this represents a targetable mechanism in disease progression, warrants direct investigation in future studies.

The parallel dysregulation of ILCs and neutrophils observed in this study, with ILCs shifting compartmentally at the CNS borders and neutrophils progressively infiltrating brain parenchyma, suggests a coordinated and self-amplifying innate immune cascade in VCID. ILCs, through their cytokine secretion at the meningeal and choroid plexus interfaces, likely create a pro-inflammatory microenvironment that facilitates neutrophil activation and subsequent parenchymal infiltration. Conversely, activated hippocampal neutrophils, through their release of IL-1β, MPO, and ROS, further amplify the neuroinflammatory signal that disrupts CNS homeostasis. This bidirectional ILC-neutrophil crosstalk, spanning from CNS borders to brain parenchyma, likely represents a key pathological mechanism driving cognitive decline in VCID.

Recent work from our group further supports the translational relevance of ILC biology in the CNS. We demonstrated that adoptively transferred ILC2s can actively traffic to and localize within CNS compartments, including in the context of glioblastoma (25). This finding underscores the migratory capacity of ILCs under pathological CNS conditions and raises the possibility that ILC2s could be harnessed as a cellular delivery platform for brain-directed immunomodulation, a strategy that may have relevance in VCID as well. Collectively, our findings identify a novel innate immune signature in VCID, characterized by compartment-specific ILC redistribution and pro-inflammatory neutrophil polarization in the meninges and choroid plexus. These immune alterations likely converge to amplify neuroinflammation, disrupt CNS homeostasis, and accelerate cognitive decline (1, 2). While the current study establishes the landscape of this innate immune dysregulation, future mechanistic studies investigating the ILC-neutrophil axis, the potential role of NETosis, and the therapeutic modulation of these pathways will be essential to translate these findings into clinical benefit for patients with VCID.

## Conclusion

This study advances our understanding of the innate immune landscape in vascular cognitive impairment and dementia by revealing, for the first time, compartment-specific dysregulation of both innate lymphoid cells and neutrophils in the meninges and choroid plexus. Our findings demonstrate that VCID drives a redistribution of ILCs, with ILC2s shifting from the meninges to the choroid plexus, alongside a pronounced pro-inflammatory N1-like neutrophil polarization in both compartments, characterized by elevated MPO and IL-1β expression and reduced anti-inflammatory markers. Critically, activated neutrophils were further shown to infiltrate the hippocampal parenchyma, revealing a spatially progressive innate immune response spanning from CNS border compartments to brain tissue. Together, these innate immune alterations converge to amplify neuroinflammation, disrupt CNS homeostasis, and accelerate cognitive decline. By recognizing that cognitive decline and neuroinflammation are not isolated processes but interconnected and compartmentalized phenomena, this work lays a foundation for future mechanistic and therapeutic studies aimed at restoring innate immune homeostasis in VCID. Targeting the ILC-neutrophil axis, through strategies such as ILC2 modulation, neutrophil polarization correction, or NETosis inhibition, may represent a promising and largely unexplored avenue for slowing or halting the progression of vascular-related cognitive decline. The next generation of research in this field may significantly reshape how we approach brain health, neuroinflammation, and aging.

## Statements

### Author contributions

All authors contributed to the study, commented on the manuscript, and approved the final version.

### Declaration of AI Assistance

AI-based software (e.g., ChatGPT) was used solely for language refinement and grammar editing; all scientific content and interpretation were entirely developed by the authors.

## Data availability

All data supporting the findings of this study are available within the paper and also are available upon request from the corresponding author.

## References

1 O’Brien JT, Thomas A. Vascular dementia. Lancet. 2015 Oct 24;386(10004):1698–706.

2 Iadecola C, Duering M, Hachinski V, Joutel A, Pendlebury ST, Schneider JA, Dichgans M. Vascular Cognitive Impairment and Dementia: JACC Scientific Expert Panel. J Am Coll Cardiol. 2019 Jul 2;73(25):3326–3344.

3 Eisenmenger LB, Peret A, Famakin BM, Spahic A, Roberts GS, Bockholt JH, Johnson KM, Paulsen JS. Vascular contributions to Alzheimer’s disease. Transl Res. 2023 Apr;254:41–53.

4 Castellani G, Croese T, Peralta Ramos JM, Schwartz M. Transforming the understanding of brain immunity. Science. 2023 Apr 7;380(6640):eabo7649.

5 Jin H, Li M, Jeong E, Castro-Martinez F, Zuker CS. A body-brain circuit that regulates body inflammatory responses. Nature. 2024 Jun;630(8017):695–703.

6 Mapunda JA, Tibar H, Regragui W, Engelhardt B. How Does the Immune System Enter the Brain? Front Immunol. 2022 Feb 22;13:805657.

7 Yuan Y, Sun J, Dong Q, Cui M. Blood-brain barrier endothelial cells in neurodegenerative diseases: Signals from the “barrier”. Front Neurosci. 2023 Feb 24;17:1047778.

8 Lin L, Chen Y, He K, Metwally S, Jha R, Capuk O, Bhuiyan MIH, Singh G, Cao G, Yin Y, Sun D. Carotid artery vascular stenosis causes the blood-CSF barrier damage and neuroinflammation. J Neuroinflammation. 2024 Sep 10;21(1):220.

9 Hu C, Zhang X, Teng T, Ma ZG, Tang QZ. Cellular Senescence in Cardiovascular Diseases: A Systematic Review. Aging Dis. 2022 Feb 1;13(1):103–128.

10 Kritsilis M, V Rizou S, Koutsoudaki PN, Evangelou K, Gorgoulis VG, Papadopoulos D. Ageing, Cellular Senescence and Neurodegenerative Disease. Int J Mol Sci. 2018 Sep 27;19(10):2937.

11 Müller L, Di Benedetto S. Inflammaging, immunosenescence, and cardiovascular aging: insights into long COVID implications. Front Cardiovasc Med. 2024 Jun 26;11:1384996.

12 Baban B, Khodadadi H, Salles ÉL, Costigliola V, Morgan JC, Hess DC, Vaibhav K, Dhandapani KM, Yu JC. Inflammaging and Cannabinoids. Ageing Res Rev. 2021 Dec;72:101487.

13 Javed K, Reddy V, Lui F. Neuroanatomy, Choroid Plexus. 2023 Jul 24. In: StatPearls [Internet]. Treasure Island (FL): StatPearls Publishing; 2024 Jan–. PMID: 30844183.

14 Ghersi-Egea JF, Strazielle N, Catala M, Silva-Vargas V, Doetsch F, Engelhardt B. Molecular anatomy and functions of the choroidal blood-cerebrospinal fluid barrier in health and disease. Acta Neuropathol. 2018 Mar;135(3):337–361.

15 Thompson D, Brissette CA, Watt JA. The choroid plexus and its role in the pathogenesis of neurological infections. Fluids Barriers CNS. 2022 Sep 10;19(1):75.

16 Lazarevic I, Soldati S, Mapunda JA, Rudolph H, Rosito M, de Oliveira AC, Enzmann G, Nishihara H, Ishikawa H, Tenenbaum T, Schroten H, Engelhardt B. The choroid plexus acts as an immune cell reservoir and brain entry site in experimental autoimmune encephalomyelitis. Fluids Barriers CNS. 2023 Jun 1;20(1):39.

17 Walker JA, Barlow JL, McKenzie AN. Innate lymphoid cells--how did we miss them? Nat Rev Immunol. 2013 Feb;13(2):75–87.

18 Srivastava RK, Sapra L, Bhardwaj A, Mishra PK, Verma B, Baig Z. Unravelling the immunobiology of innate lymphoid cells (ILCs): Implications in health and disease. Cytokine Growth Factor Rev. 2023 Dec;74:56–75.

19 Calvi M, Di Vito C, Frigo A, Trabanelli S, Jandus C, Mavilio D. Development of Human ILCs and Impact of Unconventional Cytotoxic Subsets in the Pathophysiology of Inflammatory Diseases and Cancer. Front Immunol. 2022 May 26;13:914266.

20 Baban B, Braun M, Khodadadi H, Ward A, Alverson K, Malik A, Nguyen K, Nazarian S, Hess DC, Forseen S, Post AF, Vale FL, Vender JR, Hoda MN, Akbari O, Vaibhav K, Dhandapani KM. AMPK induces regulatory innate lymphoid cells after traumatic brain injury. JCI Insight. 2021 Jan 11;6(1):e126766.

21 Huang Y, Mao K, Germain RN. Thinking differently about ILCs-Not just tissue resident and not just the same as CD4+ T-cell effectors. Immunol Rev. 2018 Nov;286(1):160–171.

22 Vivier E, Artis D, Colonna M, Diefenbach A, Di Santo JP, Eberl G, Koyasu S, Locksley RM, McKenzie ANJ, Mebius RE, Powrie F, Spits H. Innate Lymphoid Cells: 10 Years On. Cell. 2018 Aug 23;174(5):1054–1066.

23 Kveštak D, Mihalić A, Jonjić S, Brizić I. Innate lymphoid cells in neuroinflammation. Front Cell Neurosci. 2024 Feb 21;18:1364485.

24 Fung ITH, Sankar P, Zhang Y, Robison LS, Zhao X, D’Souza SS, Salinero AE, Wang Y, Qian J, Kuentzel ML, Chittur SV, Temple S, Zuloaga KL, Yang Q. Activation of group 2 innate lymphoid cells alleviates aging-associated cognitive decline. J Exp Med. 2020 Apr 6;217(4):e20190915.

25 Wang LP, Bhandari B, Emami Naeini S, Yu JC, Arbab AS, Young N, Lopes Salles É, Baban B. Adoptive transfer of ILC2s reveals tumor homing in glioblastoma: a proof-of-concept study. Front Oncol. 2026;16:1776061.

26 Shafqat A, Noor Eddin A, Adi G, Al-Rimawi M, Abdul Rab S, Abu-Shaar M, Adi K, Alkattan K, Yaqinuddin A. Neutrophil extracellular traps in central nervous system pathologies: A mini review. Front Med (Lausanne). 2023 Feb 17;10:1083242.

27 Santos-Lima B, Pietronigro EC, Terrabuio E, Zenaro E, Constantin G. The role of neutrophils in the dysfunction of central nervous system barriers. Front Aging Neurosci. 2022 Aug 11;14:965169.

28 Mihaila AC, Ciortan L, Macarie RD, Vadana M, Cecoltan S, Preda MB, Hudita A, Gan AM, Jakobsson G, Tucureanu MM, Barbu E, Balanescu S, Simionescu M, Schiopu A, Butoi E. Transcriptional Profiling and Functional Analysis of N1/N2 Neutrophils Reveal an Immunomodulatory Effect of S100A9-Blockade on the Pro-Inflammatory N1 Subpopulation. Front Immunol. 2021 Aug 10;12:708770.

29 Vaibhav K, Braun M, Alverson K, Khodadadi H, Kutiyanawalla A, Ward A, Banerjee C, Sparks T, Malik A, Rashid MH, Khan MB, Waters MF, Hess DC, Arbab AS, Vender JR, Hoda N, Baban B, Dhandapani KM. Neutrophil extracellular traps exacerbate neurological deficits after traumatic brain injury. Sci Adv. 2020;6(22):eaax8847.

30 Khan MB, Alam H, Siddiqui S, Shaikh MF, Sharma A, Rehman A, Baban B, Arbab AS, Hess DC. Exercise Improves Cerebral Blood Flow and Functional Outcomes in an Experimental Mouse Model of Vascular Cognitive Impairment and Dementia (VCID). Transl Stroke Res. 2024 Apr;15(2):446–461.

31 Khodadadi H, Salles ÉL, Jarrahi A, Costigliola V, Khan MB, Yu JC, Morgan JC, Hess DC, Vaibhav K, Dhandapani KM, Baban B. Cannabidiol Ameliorates Cognitive Function via Regulation of IL-33 and TREM2 Upregulation in a Murine Model of Alzheimer’s Disease. J Alzheimers Dis. 2021;80(3):973–977.

32 Khodadadi H, Salles ÉL, Naeini SE, Bhandari B, Rogers HM, Gouron J, Meeks W, Terry AV Jr, Pillai A, Yu JC, Morgan JC, Vaibhav K, Hess DC, Dhandapani KM, Wang LP, Baban B. Boosting Acetylcholine Signaling by Cannabidiol in a Murine Model of Alzheimer’s Disease. Int J Mol Sci. 2024 Nov 1;25(21):11764.

33 Tian Z, Ji X, Liu J. Neuroinflammation in Vascular Cognitive Impairment and Dementia: Current Evidence, Advances, and Prospects. Int J Mol Sci. 2022 Jun 2;23(11):6224. doi: 10.3390/ijms23116224. PMID: 35682903; PMCID: PMC9181710.

34 Smyth LCD, Murray HC, Hill M, van Leeuwen E, Highet B, Magon NJ, Osanlouy M, Mathiesen SN, Mockett B, Singh-Bains MK, Morris VK, Clarkson AN, Curtis MA, Abraham WC, Hughes SM, Faull RLM, Kettle AJ, Dragunow M, Hampton MB. Neutrophil-vascular interactions drive myeloperoxidase accumulation in the brain in Alzheimer’s disease. Acta Neuropathol Commun. 2022 Mar 24;10(1):38.

